# A cell non-autonomous mechanism of yeast chronological aging regulated by caloric restriction and one-carbon metabolism

**DOI:** 10.1101/2020.07.13.200493

**Authors:** Elisa Enriquez-Hesles, Daniel L. Smith, Nazif Maqani, Margaret B. Wierman, Matthew Sutcliffe, Ryan D. Fine, Agata Kalita, Sean M. Santos, Michael J. Muehlbauer, James R. Bain, Kevin A. Janes, John L. Hartman, Matthew D. Hirschey, Jeffrey S. Smith

**Affiliations:** Department of Biochemistry and Molecular Genetics, University of Virginia School of Medicine, Charlottesville, VA; Department of Biomedical Engineering, University of Virginia School of Medicine, Charlottesville, VA; Department of Nutrition Science, Nutrition and Obesity Research Center, Nathan Shock Center of Excellence in the Basic Biology of Aging, University of Alabama at Birmingham, Birmingham, AL; Department of Genetics, Nutrition and Obesity Research Center, Nathan Shock Center of Excellence in the Basic Biology of Aging, University of Alabama at Birmingham, Birmingham, AL; Department of Medicine, Duke Molecular Physiology Institute, Duke University, Durham, NC

## Abstract

Caloric restriction (CR) improves healthspan and lifespan of organisms ranging from yeast to mammals. Understanding the mechanisms involved will uncover future interventions for aging associated diseases. In budding yeast, *Saccharomyces cerevisiae*, CR is commonly defined by reduced glucose in the growth medium, which extends both replicative and chronological lifespan (CLS). We found that conditioned media collected from stationary phase CR cultures extended CLS when supplemented into non-restricted (NR) cultures, suggesting a potential cell non-autonomous mechanism of CR-induced lifespan regulation. Chromatography and untargeted metabolomics of the conditioned media, as well as transcriptional responses associated with the longevity effect, pointed to specific amino acids enriched in the CR conditioned media (CRCM) as functional molecules, with L-serine being a particularly strong candidate. Indeed, supplementing L-serine into NR cultures extended CLS through a mechanism dependent on the one-carbon metabolism pathway, thus implicating this conserved and central metabolic hub in lifespan regulation.

## Introduction

Caloric restriction (CR) extends lifespan in a wide variety of model organisms ranging from the budding yeast, *Saccharomyces cerevisiae*, to non-human primates, implying that conserved cellular processes and pathways must mediate the beneficial effects, or somehow be impacted by the dietary regimen (1). Indeed, conserved processes including autophagy, TOR signaling, and AMPK signaling have each been implicated in regulating aging in most models of CR (2). In the yeast system, CR is typically characterized by reducing the initial glucose concentration in growth medium from 2% (non-restricted; NR) to 0.5% or lower, or reducing overall amino acids (3, 4). Glucose restriction robustly extends both yeast replicative lifespan (RLS) and chronological lifespan (CLS) (3–6), the latter of which is defined by the number of days that non-dividing cells maintain proliferative capacity in liquid culture after entering stationary phase, quantified upon transfer to fresh nutrient media (7, 8). As glucose becomes limiting toward the end of exponential growth, cells switch from fermentative to mitochondrial-driven oxidative metabolism of the ethanol and organic acids produced during fermentation. This ‘diauxic shift’ is accompanied by dramatic changes in transcription, translation, and metabolic profiles that facilitate slower cell growth using non-fermentable carbon sources (9, 10), ultimately leading to cell cycle exit and quiescence. CLS largely hinges on an adaptive response to nutrient depletion, consisting of cell cycle exit (G0), called quiescence (11, 12). Yeast CLS is therefore considered a model for the aging of quiescent stem cells, or post-mitotic cells like neurons or muscle fiber cells (13–15).

In yeast CLS assays, glucose concentration in media (2% or 0.5%) is usually specified at the time of culture inoculation. Measurements of cell viability (colony forming units) are then initiated 3 or 4 days later, after nutrients are depleted and cell proliferation ceases. Therefore, understanding intracellular and extracellular responses underlying the adaptive transition to quiescence, and how CR influences them, are of central importance. CR enhances several processes that occur during the diauxic shift, including Snf1 (AMPK) signaling (16), mitochondrial respiration and ATP production (6, 17–19), accumulation of the storage carbohydrate trehalose (20), and improved G1 cell cycle arrest (21). Several other conserved genetic and environmental manipulations, such as inhibition of TOR signaling (22) and methionine restriction (23, 24), also extend CLS. As unicellular organisms, the impact of such conditions on longevity is primarily expected to occur through cell autonomous mechanisms such as changes in gene expression and metabolism. However, regulation of CLS by genetic and environmental manipulations is also linked with cell non-autonomous effects (25, 26). Low pH and high acetic acid concentrations are associated with apoptosis and reduced CLS, which can be suppressed by CR, TOR inhibition, buffering against media acidification, or even transferring cells to water after stationary phase (25, 27). Acetic acid stress also activates nutrient sensing growth pathways that lead to elevated superoxide (28).

Longevity-associated cell non-autonomous mechanisms are classically described from rodent models, where circulating extracellular factors have been identified from heterochronic parabiosis experiments (29). For example, mesencephalic astrocyte-derived neurotrophic factor (MANF) from younger mice protects against liver damage in the older mice, and its overexpression extends lifespan in *Drosophila* (30). Furthermore, some factors that act in a cell autonomous manner, such as the insulin-like signaling transcription factor FOXO, can impact organismal longevity via cell non-autonomous mechanisms (31, 32), raising the possibility that such processes are more widespread than previously thought.

Despite being single cell organisms, budding yeast utilize proteins and metabolites for cell-cell communication associated with mating, differentiation and sporulation. Recognition of opposite haploid mating types (a-cells or α-cells) occurs via the extracellular pheromone peptides a-factor and α-factor (33), whereas pseudohyphal growth in dense cultures or colonies is mediated by quorum sensing via the amino acid derived aromatic alcohols, tryptophol and phenylethanol (34). Chronological aging of *S. cerevisiae*, which occurs in densely crowded cultures and is highly sensitive to gene-nutrient interactions (35), would also seem subject to cell non-autonomous mechanisms. Indeed, unidentified high molecular weight (>5,000 Da) extracellular factors from old stationary phase cultures have been implicated in stimulating survival of other old cells (36). Similarly, our lab observed that conditioned media from glucose-restricted stationary phase cultures extended CLS when supplemented into non-restricted cultures (37), suggesting the presence of one or more extracellular proteins, peptides, or metabolites that contribute to lifespan regulation. Determining the identity of such factors would provide new insights about CR mechanisms. Here, we have utilized a combination of chromatography, metabolomics, and targeted mass spectrometry to identify functional candidate factors that were more abundant in CR conditioned media (CRCM) than in NR-conditioned media (NRCM). Longevity activity was traced to multiple amino acids that extend CLS when supplemented into NR cultures. We focused further analysis on L-serine, and in the process, implicated the one-carbon metabolism pathway in CLS regulation.

## RESULTS

### Conditioned media from CR stationary phase cultures contains longevity factors

It is well established that CR in the form of glucose restriction extends CLS of yeast cells (5, 6). Media swap experiments indicated at least part of this effect was due to differences in the growth medium (25, 38). For example, when 5-day old NR and CR stationary phase cultures of strain BY4741 were pelleted and the conditioned media exchanged (Figure 1A), NR-grown cells displayed increased CLS when transferred into the CRCM (Figure 1B, left panel), while the long CLS of CR-grown cells was lost when transferred into NRCM (Figure 1B, right panel). This reciprocal effect on CLS was previously ascribed to differences in pH and organic acids such as acetic acid (25). To determine if additional factors in the conditioned media were involved in CLS extension, we concentrated the NRCM and CRCM from stationary phase cultures using a Rotavap apparatus, supplemented each concentrate into non-restricted cultures at the time of inoculation (1% vol/vol), and then assayed CLS using a quantitative microcolony viability assay (see (16) and Experimental Procedures). The CRCM concentrate significantly extended mean CLS, while the NRCM did not (Figure 1C and D). We next confirmed the difference between CRCM and NRCM by titrating in higher amounts of each concentrate (2%, 5%, and 10% vol/vol) and utilizing an independent high-throughput CLS assay in a 384-well format that allowed for 96 replicates per condition (Figure S1A). With this system, improved CLS is indicated by a lower L parameter on the y-axis, representing the time it takes a small aliquot spotted onto a fresh YPD plate to reach ½ maximum growth density (35, 39). More viable cells equal less time to ½ max growth. Compared to the controls (dH_2_O and NRCM supplements), the CRCM supplement again showed concentration-dependent CLS extension (Figure S1A), lower L parameter). NRCM only had minor effects at the higher (5% and 10%) supplement levels. We therefore compared the CR and NRCM concentrates at 1% and 2% (vol/vol) in the quantitative microcolony CLS assay (Figure S1B), C, and E). Weak CLS extension was observed even with the NRCM concentrate at 2% (vol/vol), suggesting the existence of active compounds in both types of conditioned media, with higher levels in the CRCM. Since BY4741 is auxotrophic for histidine, leucine, methionine, and uracil, we confirmed that CRCM isolated from BY4741 also extended the CLS of a prototrophic strain, FY4 (Figure S1D and F).

**Figure 1.**
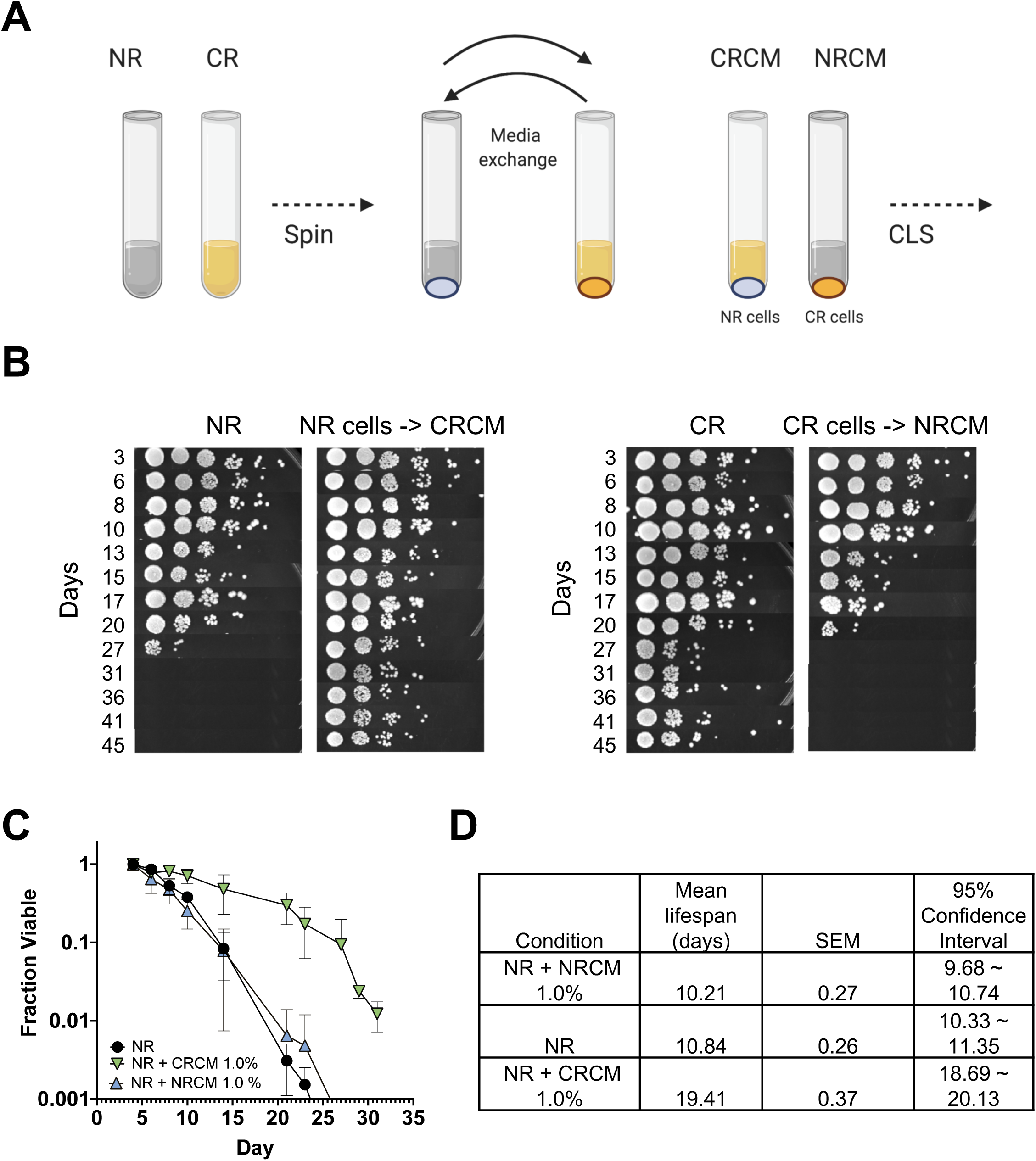
Longevity factors present in conditioned media from calorie restricted treated yeast cultures. **(A)** Schematic of basic media exchange experiment where WT lab strain BY4741 was grown to stationary phase in SC media containing 2% (NR) or 0.5% (CR) glucose. The cells were pelleted by centrifugation and the conditioned media then filtered and exchanged, such that NR-grown cells will age in CR conditioned media (CRCM) and CR-grown cells will age in NR conditioned media (NRCM). **(B)** Qualitative spot test assay tracking cell over time, starting at the time of media exchange (day 3). The NR and CR controls (left panels) represent samples where the conditioned media was filtered but not exchanged. **(C)** Quantitative chronological life span (CLS) assay. Concentrated CRCM and NRCM was supplemented at 2% (vol/vol) into NR cultures at time of inoculation. To measure the fraction of viable cells over time, micro-colony forming units were counted after 18 hours of regrowth after spotting onto YPD plates. Error bars indicate standard deviations (n=3). **(D)** Mean lifespan in days was calculated using Online Application for Survival Analysis Software (OASIS 2).

### Fractionation of CRCM isolates CLS factor activities separate from acetic acid

An earlier study concluded that chronologically aged yeast cells release large (>5 kD) heat-stable compounds into the media that improve viability of other cells in the population (36). To determine if CRCM contained such factors, we treated it with Proteinase K, DNase I, RNase A, phenol/chloroform, autoclaving, or freezing, but none of these had any effect on CLS extension (data not shown). Instead, CRCM molecular activity was found to be smaller than 5,000 daltons, as the fraction passing through an Amicon Ultra-4 centrifugal filter unit (5,000 MW cutoff) had CLS extending activity equivalent to the starting material (Figure 2A). This result demonstrated that the CR-induced longevity factor(s) described here were different from the previously described higher molecular weight factors (36).

**Figure 2.**
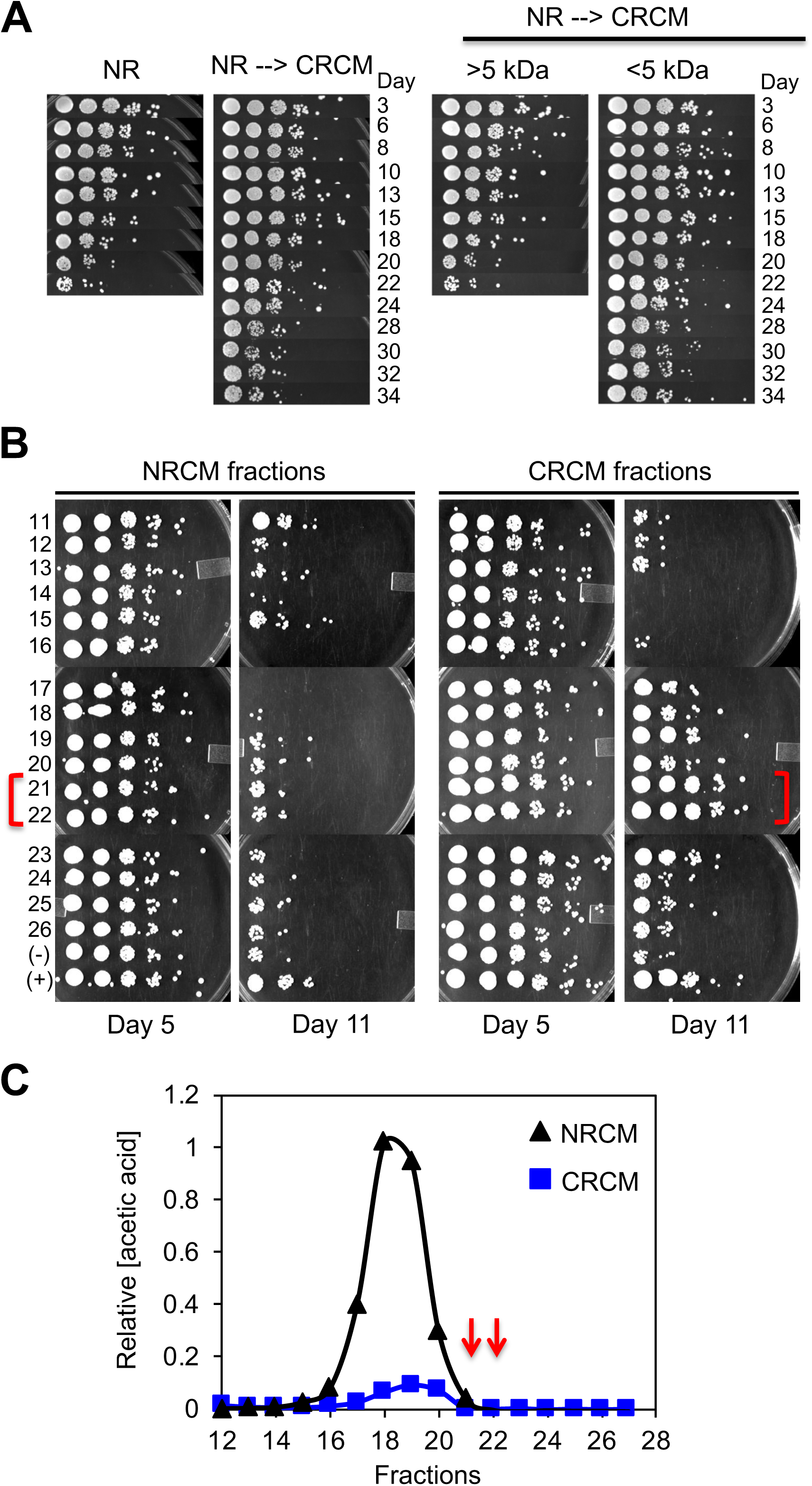
Chromatographic sizing and separation of longevity factor activity in CRCM. **(A)** Left panel: Qualitative CLS assay showing improved viability when supplementing CRCM into non-restricted BY4741 cultures. Right panel: CRCM was first separated into high MW (>5 kDa) or low MW (<5 kDa) fractions using an Amicon Ultra-4 centrifugal filter unit, then supplemented into non-restricted BY4741 cultures. Longevity activity was retained in the low MW fraction. **(B)** Size exclusion chromatography of NRCM and CRCM was performed using a Sephadex G-10 column (700 dalton MW cutoff). Fractions were added to non-restricted BY4741 cultures at the time of inoculation, and viability tracked over time with qualitative spot assays. The effects of fractions 11 to 26 are shown for days 5 and 11. The longevity peak fractions for CRCM are bracketed in red. **(C)** Relative acetic acid concentration was measured in the NRCM and CRCM fractions. Red arrows indicate the fractions with longevity activity in CRCM.

To better separate CLS-modifying activities in the conditioned media, we concentrated 150 ml of NRCM or CRCM down to a final volume of 2.5 ml, removed any precipitates by centrifugation, and then fractionated the soluble material through a Sephadex G-10 column, which has a size exclusion limit of ∼700 Da. Fractions were then added to NR cultures of BY4741 at a 1:5 ratio and CLS extension detected using a qualitative spot test assay (Figure 2B). At day 11, there was a clear peak of improved viability at fractions 21 and 22 for the CRCM, suggesting the active compounds were smaller than 700 Da (Figure 2B). We considered the possibility that high levels of acetic acid in the NRCM could potentially mask longevity activity in the fractions. However, acetic acid peaked at fractions 18-19 in these columns, distinct from the CRCM longevity peak at fractions 21-22 (Figure 2C, red arrows). Instead, the low day 11 viability with NRCM fractions 16-18 was potentially due to elevated acetic acid (Figure 2B and C). Based on this size exclusion chromatography and the resistance to various treatments such as heat, phenol extraction, nuclease digestion, etc., we concluded that the longevity factor(s) in CRCM were small water-soluble compounds separable from acetic acid.

### CRCM-enriched metabolites and induced genes indicate amino acids modulate CLS

To identify candidate small molecule longevity factors in the CRCM, we utilized a comparative metabolomics approach to generate metabolite profiles for the CR and NR conditioned media (Table S1), reasoning that differential abundance of extracellular metabolites could be a source of CRCM longevity factors (Figure 3A). Enrichment and pathway analysis of metabolites more abundant in the CRCM compared to NRCM was performed using MetaboAnalyst (40). Most of the significantly enriched pathways were related to amino acids, including L-alanine, L-aspartate, and L-glutamate metabolism, as well as L-glycine, L-serine, and L-threonine biosynthesis, both of which had the two highest pathway impact scores (Figure 3B), a combined representation of centrality and pathway enrichment (40).

**Figure 3.**
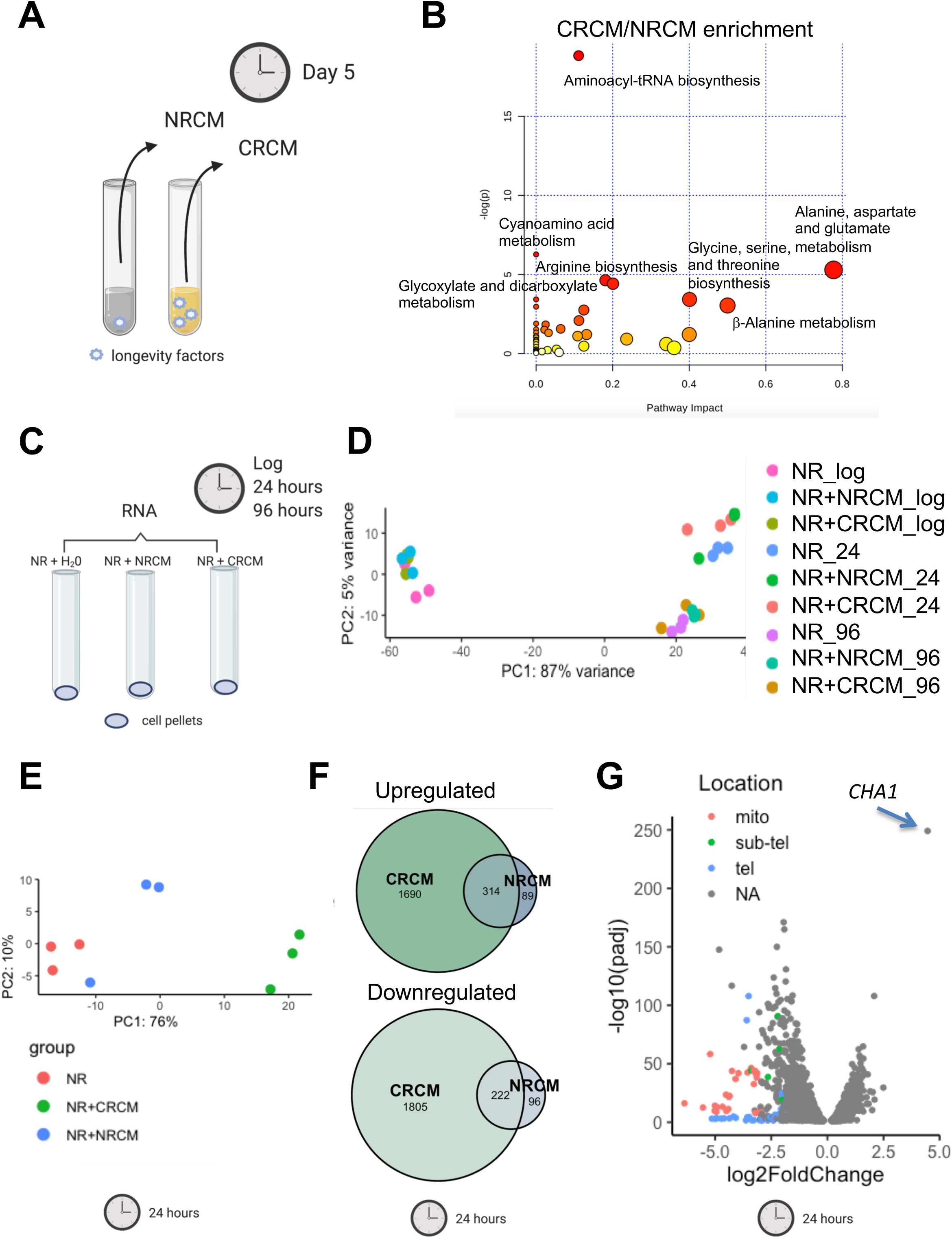
Metabolomic and RNA-seq analysis point toward amino acid metabolism. **(A)** Schematic of conditioned medium collection used for untargeted metabolomics analysis after five days of SC cultures grown with 2.0% (NR) or 0.5% glucose (CR). **(B)** MetaboAnalyst software was used for Enrichment and Pathway Analysis of metabolites with a CR/NR ratio greater then 1.0. The names of KEGG pathways with p<0.05 are highlighted. Pathway impact is a measure that considers the centrality of a metabolite in the pathway. Circle size is proportional to pathway impact value and red color indicates more significant changes. **(C)** Schematic of cell conditions (NR, NR + 2.0 % NRCM, and NR + 2.0% CRCM) collected for RNA-seq analysis at log phase, 24 hours, and 96 hours. **(D)** Principal component analysis of RNA-seq samples at log, 24, and 96 hour conditions of NR, NR+ NRCM, and NR+ CRCM. **(E)** Principal component analysis of RNA-seq samples collected at 24 hours. **(F)** Venn diagram of differentially expressed genes (up or down; FDR<0.05) for NRCM- or CRCM-supplemented samples, as compared to the NR + H_2_O control at 24 hours. **(G)** Volcano plot displaying differential expressed genes between the NR + CRCM and NR + H_2_O samples at 24 hours. The y-axis indicates the p-adjusted value and x-axis the log2 fold change. Red, green, and blue denotes genes located in the mitochondrial genome, sub-telomeric, or telomeric regions, respectively. The most upregulated gene *CHA1* is highlighted by an arrow.

To gain additional insights to candidate factors that were potentially functional, we also performed transcriptomics analysis on BY4741 cells grown in non-restricted SC media supplemented with CRCM, NRCM, or water as a control (Figure 3C), with a goal of identifying physiological responses linked to specific metabolites. Following inoculation into these conditions, we harvested cells at log phase, 24 hrs (late diauxic shift), and 96 hrs (stationary phase), then performed RNA-seq on isolated mRNAs. Principal component analysis (PCA) indicated the major variance within the data was the time points (Figure 3D), consistent with the massive transcriptional changes that occur during the transition into stationary phase (9, 10). In early log phase cells, there were no significantly upregulated or downregulated genes in the CRCM- or NRCM-supplemented samples as compared to the H_2_O-supplemented control (FDR <0.05), consistent with earlier microarray analysis showing that CR (0.5% glucose) had little effect on gene expression during early log phase (37). At the 24 hr timepoint, however, CRCM-supplemented samples diverged from the NRCM- and H_2_O-supplemented controls in a PCA plot (Figure 3E), and showed many more differentially regulated genes than the NRCM-treated samples (Figure 3F). At the 96 hr timepoint, gene expression for the NRCM samples also clearly differentiated from the H_2_O-supplemented control (Figure S2A), though there were still a large number of genes exclusively altered in the CRCM samples (Figure S2B). The top GO term for CRCM-upregulated genes at 96 hr was *α*-amino acid catabolic process (Table S2), consistent with the metabolomics results indicating amino acid metabolism. Interestingly, there were a number of telomeric and sub-telomeric ORFs that were more tightly repressed in the CRCM-treated cells compared to the NR control at 24 and 96 hr (Figures S2C, 3G, and Tables S4, S5), suggesting that the general transcriptional repression associated with chromatin condensation in quiescent cells may be enhanced by supplementing with CRCM (41, 42). At 24 hrs, the top GO terms for CRCM-upregulated genes were related to mitochondrial function and respiration, consistent with a more robust metabolic transition during the diauxic shift (Table S3).

Furthermore, the *YCL064C* (*CHA1*) gene, which is adjacent to the heterochromatic *HML* locus, clearly stood out as the most significantly upregulated (Figure 3G and Table S4). *CHA1* encodes a predominantly mitochondrial L-serine (L-threonine) deaminase that catabolizes these amino acids as nitrogen sources, and in the case of L-serine, for direct production of pyruvate (43, 44).

It is strongly upregulated by exogenous L-serine or L-threonine added to the growth medium (43, 45), suggesting that L-serine and possibly L-threonine in the CRCM could be producing an especially strong physiological response related to CLS extension. Together, the extracellular metabolite analysis and effects on gene expression during the diauxic shift and stationary phase pointed toward amino acids, especially L-serine, as candidate extracellular CLS factors mediating the CR effect on CLS.

### Amino Acids are depleted from NR conditioned stationary phase media

We next profiled all 20 standard amino acids from BY4741 CRCM and NRCM concentrates, as well as unconditioned (fresh) SC media that was concentrated in the same manner (Figure 4A). All but 6 amino acids (alanine, cysteine, glutamine, glycine, proline, and valine) were significantly depleted to varying degrees in NRCM concentrate, relative to unconditioned SC concentrate. CR strongly attenuated the depletion, indicating that amino acid levels were generally higher in CRCM than NRCM. L-Serine is an excellent example of this relationship (Figure 4A). The CR/NR abundance ratios for lysine, asparagine, and serine were each 10-fold or higher in the CRCM (Figure 4B), but still less than the level in unconditioned SC concentrate (Figure 4A). Notably, the branched chain amino acids leucine and valine were significantly more abundant in CRCM concentrate than SC (Figures 4A, B and S3), with isoleucine trending in the same direction, suggesting that biosynthesis and release of these amino acids was induced by CR. This effect was lost in the prototrophic FY4 strain, suggesting the *leu2Δ* mutation in BY4741 could be a contributing factor. Otherwise, the pattern of CR rescuing amino acid depletion from the media was recapitulated with FY4, though stronger NR depletion rendered CR/NR ratios more extreme (Figure 4C and D). Accordingly, the CRCM concentrate isolated from prototrophic FY4 stationary phase cultures was also effective at extending FY4 CLS (Figure 4E and F). Based on these results we hypothesized that CR was contributing to CLS extension by altering amino acid metabolism in such a way that prevented depletion from the media. The higher amino acid levels in CRCM concentrate could therefore explain why it was more effective than NRCM concentrate at extending CLS. Consistent with this interpretation, supplementing the concentrated unconditioned SC media into NR cultures also extended CLS (Figure 4G and H).

**Figure 4.**
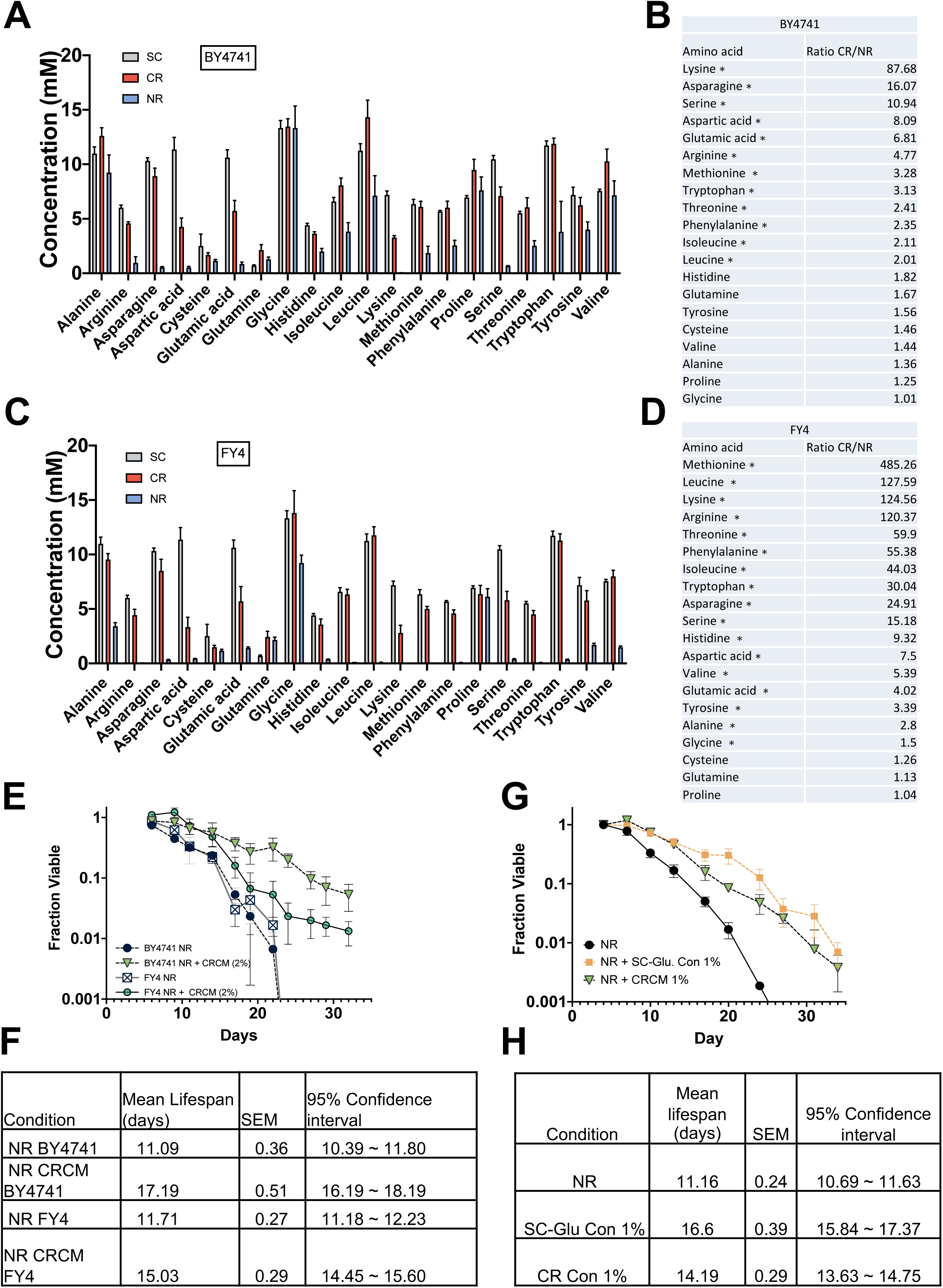
CR conditioned media is enriched for multiple amino acids. **(A)** Quantification of amino acids in NRCM and CRCM from stationary phase BY4741 cultures, or the starting unconditioned SC media (concentrated 10-fold). Amino acids were separated with a ZipChip and quantified by mass spectrometry. Error bars indicate standard deviation (n=3). **(B)** Amino acid abundance ratios between BY4741 CRCM and NRCM are sorted from highest to lowest. **(C)** Quantification of amino acids in concentrated NRCM and CRCM from stationary phase FY4 cultures, or unconditioned SC media. **(D)** Amino acid abundance ratios between FY4 CRCM and NRCM sorted from highest to lowest. Significant differences in panels B and D are indicated (\**p*≤0.05) using a student t-test. **(E)** Quantitative CLS of BY4741 and FY4, each supplemented with CRCM isolated from BY4741 at 2% (vol/vol). **(F)** Mean CLS statistics from panel E calculated using OASIS 2. **(G)** Quantitative CLS of BY4741 supplemented with CRCM or unconditioned SC concentrate at 1% (vol/vol). **(H)** Mean CLS statistics from panel G calculated using OASIS 2.

### Supplementation of specific amino acids is sufficient to extend CLS

Since most amino acids were depleted from stationary phase NR cultures, we reasoned that one or more of them were critical for maintaining longevity. We initially focused on L-serine because the biosynthesis gene *SER1* was previously identified as a strong quantitative trait locus (QTL) for CLS in the BY4741 background (46), and *CHA1* expression was strongly induced by CRCM supplementation during the diauxic shift (Figure 3G). The concentration of serine in our standard SC media is 1 mM (38, 47), so we tested the effect of supplementing an additional 1 mM or 5 mM L-serine into NR cultures at the time of inoculation. 5 mM L-serine significantly extended CLS, while 1 mM did not (Figure 5A and B). To confirm the L-serine effect was not specific to SC media, we also tested for CLS extension in a custom synthetic growth medium (HL) designed to support longevity that does not have ammonium sulfate as a nitrogen source (35, 48). BY4741 had significantly longer CLS in non-restricted HL medium compared to SC medium, and 5 mM L-serine further extended it (Figure S4A and B). We next tested whether other amino acids could extend CLS at 5mM (Figure 5C and D). Some amino acids did extend CLS, but not always as predicted based on abundance in the conditioned media. For example, L-asparagine had a similar depletion/enrichment profile as L-serine (Figure 4A), but did not extend CLS when added back (Figure 5C and D). We also tested supplementation with 5 mM L-glycine, a component of one-carbon metabolism that can be derived from L-serine, but was not depleted from the NR media (Figure 4A). L-glycine had no effect on CLS at this concentration (Figure 5C and D). L-cysteine supplementation at 5 mM dramatically slowed cell growth such that CFUs were increasing until day 10, after which the decline in CLS was parallel to the NR control, suggesting a delay rather than a true extension of survival (Figure 5C and D).

**Figure 5.**
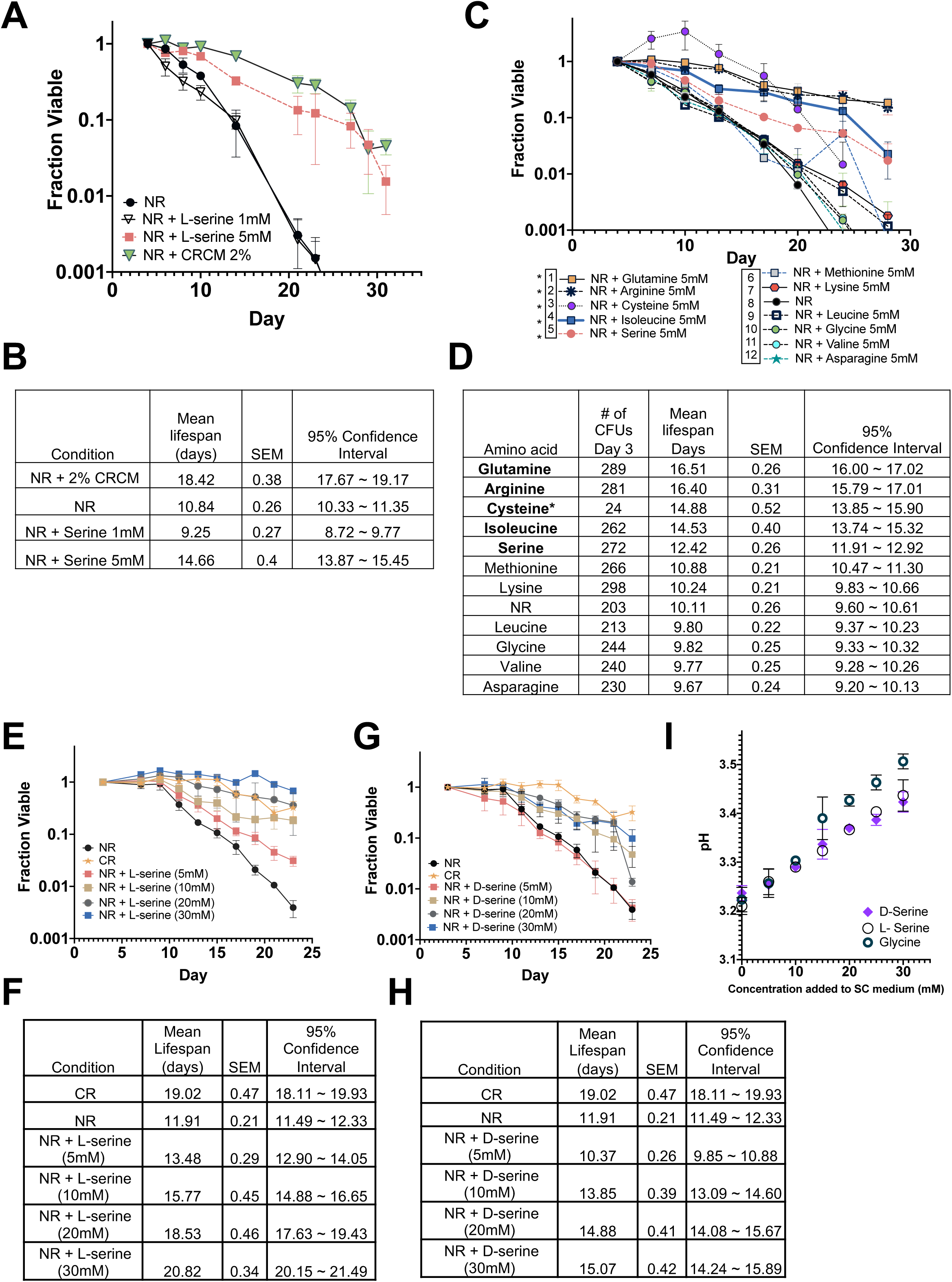
L-Serine supplementation extends lifespan of NR cultures. **(A)** CLS of non-restricted BY4741 supplemented with 2% CRCM, 1 mM L-serine, or 5 mM L-serine. **(B)** Mean CLS statistics from panel A calculated using OASIS 2. **(C)** CLS of non-restricted BY4741 supplemented with 5 mM of each indicated amino acid. *Amino acids 1-5 significantly extended lifespan compared to the NR control (sample 8) **(D)** Mean CLS statistics from panel C calculated using OASIS 2. **(E)** CLS of non-restricted BY4741 supplemented with increasing concentrations of L-serine. CR indicates the glucose-restricted control samples. **(F)** Mean CLS statistics from panel E calculated using OASIS 2. **(G**) CLS of BY4741 supplemented with increasing concentrations of D-serine. CR indicates the glucose-restricted control. **(H)** Mean CLS statistics from panel G calculated using OASIS 2. **(I)** pH measurements of SC (2% glucose) medium after supplementation with L-glycine, L-serine, or D-serine at the indicated concentrations before cell growth.

### L-serine extends CLS through catabolic and non-catabolic pathways

An earlier study of cellular response to L-serine supplementation found that its uptake was linear with increasing extracellular concentrations up to at least 100 mM (45), suggesting to us that L-serine concentrations higher than 5mM may induce stronger CLS extension. To test this idea, we supplemented NR cultures of BY4741 with 5, 10, 20, or 30 mM L-serine and observed progressively improved longevity with increasing concentration (Figure 5E and F). Survival with 30 mM L-serine was even slightly better than the CR control, showing minimal loss of viability during the experiment. A similar positive correlation between L-serine concentration and CLS was observed with FY4 (Figure S4C and D). To further examine whether improved CLS might result from catabolism of L-serine, we first supplemented NR cultures of BY4741 with the presumably non-utilized stereoisomer D-serine. We confirmed its inactivity by showing that 5 or 30 mM D-serine could not rescue the partial L-serine auxotrophic phenotype of a *ser2Δ* mutant in SC-serine media (Figure S4E-H). D-serine supplementation into BY4741 NR cultures had no effect on CLS at 5 mM (Figure 5G and H). It indistinguishably extended CLS at 10, 20, and 30 mM concentrations, but to a lesser extent than CR or the equivalent concentration of L-serine (Figure 5G and H). Based on these results we hypothesized that L-serine catabolism may be important for supporting CLS under NR conditions up to 5 or 10 mM, but additional non-catalytic mechanisms are involved at higher L-serine concentrations. An independent report concluded that exogenous amino acids, including L-serine, support CLS by preventing hyperacidification of the media (49). In our system, however, L-serine and D-serine had exactly the same pH buffering capacity on SC media (Figure 5I), even though L-serine was more effective at extending CLS. Moreover, L-glycine showed better pH buffering than L-serine (Figure 5I), but was not as effective at extending CLS (Figure 5C). We conclude that L-serine catabolism and pH buffering contribute to CLS extension through distinct mechanisms.

### L-serine extends CLS through the one-carbon metabolism pathway

CR buffers the acidification of conditioned media by promoting consumption of acetate and acetic acid via Snf1/AMPK-dependent activation of gluconeogenesis and glyoxylate cycle gene transcription (16, 50). Since L-serine accumulated in the CR conditioned media (Figure 4A), we next tested whether supplementing L-serine into NR media also promoted acetic acid consumption. As shown in Figure 6A, adding 5, 10, or 20 mM L-serine did not reduce acetic acid levels in the NR media, implying that serine extends CLS through a mechanism different from CR. Indeed, L-serine further extended CLS when added to CR cultures, again supporting the idea of independent mechanisms (Figure 6B and C).

**Figure 6.**
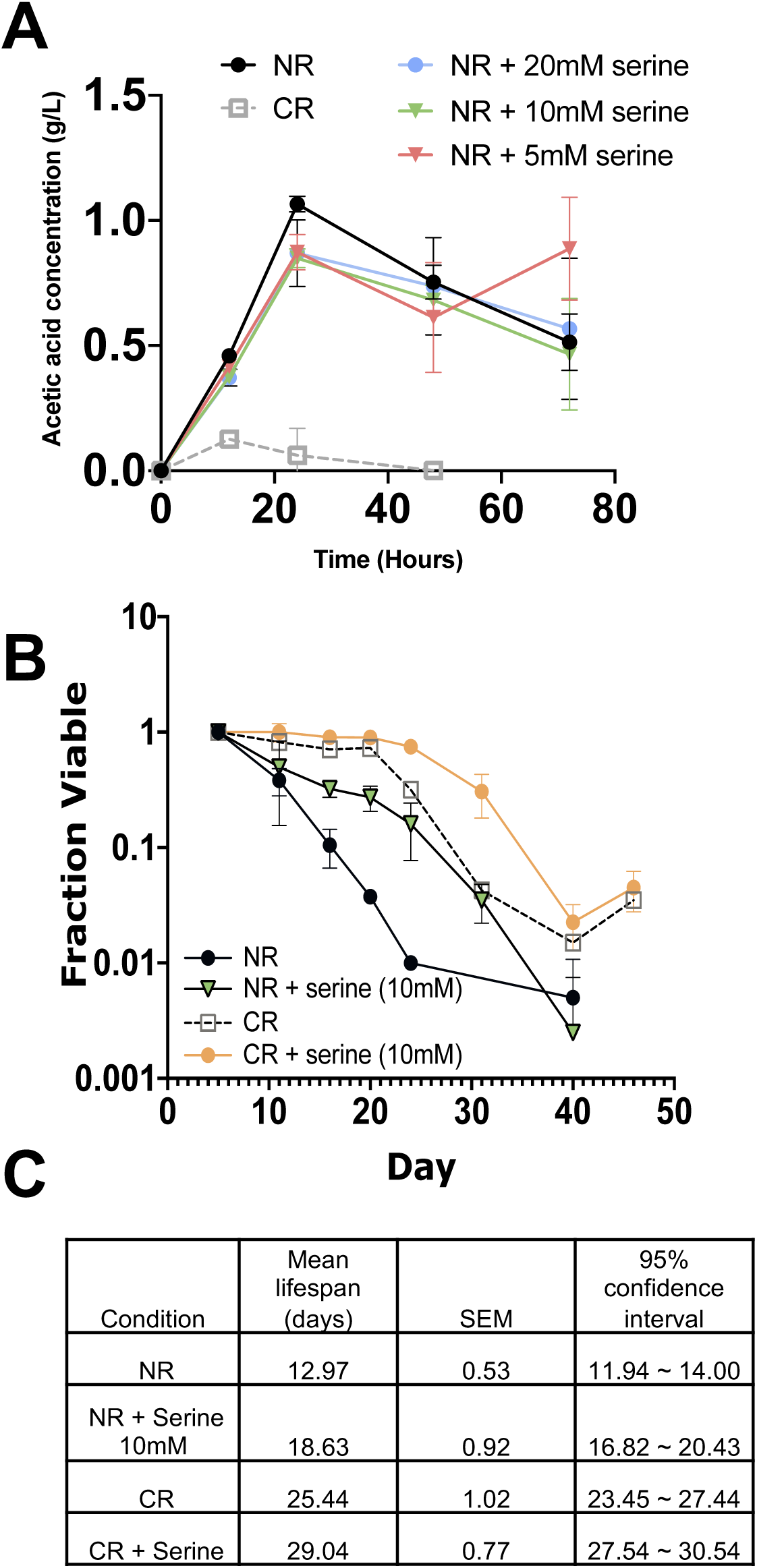
CR and serine supplementation mediate longevity by independent mechanisms. Acetic acid content in the media of CR, NR, NR + L-serine (5, 10, 20 mM) cultures was measured over the first 72 hours of the aging assays (mean ± SD, n=3). **(B)** CLS of BY4741 growing in the NR or CR media condition supplemented with 10 mM L-serine. **(C)** Mean CLS statistics from panel B calculated using OASIS 2.

Since L-serine is the predominant donor of carbon units to folate in one-carbon metabolism, we hypothesized its depletion would constrain this route of utilization in supporting CLS, which could explain extension by exogenous L-serine. If true, then mutations that impair one-carbon metabolism should attenuate the effect. The one-carbon metabolism pathway for *S. cerevisiae* is depicted in Figure 7A, including serine hydroxymethyltransferases (SHMTs), Shm1 and Shm2, that interconvert L-serine and L-glycine in the mitochondria or cytoplasm, respectively. Shm2 is the major isozyme for converting L-serine to L-glycine and one-carbon units on tetrahydrofolate, whereas Shm1 is the predominant isozyme for the reverse reaction, though their relative activities are strongly influenced by nutrient availability and growth conditions (51). We therefore supplemented NR cultures of *shm1Δ* or *shm2Δ* mutants from the YKO collection with 5 mM L-serine to observe any effects on CLS. Without serine supplementation, the *shm1Δ* mutant showed moderate extension of mean CLS when compared to WT NR cultures (Figure 7B and S5), while the *shm2Δ* mutant only showed modest improvements in survival at the later time points (Figure 7C and S5). Importantly, both deletions prevented further CLS extension induced by 5 mM L-serine, but did not attenuate the strong positive lifespan effect of CR. A similar result was obtained for a strain lacking *MTD1* (Figure 7D and S5), which encodes a cytoplasmic NAD^+^-dependent 5,10-methylenetetrahydrofolate dehydrogenase. Lastly, we followed up on a recent report that Fungal Sideroflexin-1 (Fsf1) is a functional homolog of the human Sideroflexin-1 protein SFXN1, which was discovered through a CRISPR screen to be the mitochondrial serine transporter for one-carbon metabolism (52, 53). Ectopic expression of yeast *FSF1* in an SFXN1 mutant cell line rescued its defects in mitochondrial serine transport and *de novo* purine synthesis (52), suggesting that Fsf1 could have the same function in yeast mitochondria, where it is localized (54). We therefore tested whether cells deleted for *FSF1* would show the same lack of responsiveness to L-serine supplementation, and also tested higher concentrations of L-serine. There was no indication of CLS extension for the *fsf1Δ* mutant under the NR condition (Figure 7E and F). However, the effects of 5 mM and 10 mM L-serine on CLS were blocked or attenuated, respectively (Figure 7E and F). CLS was still strongly extended by 30 mM L-serine, most likely due to the pH buffering effect (Figure 7E). Based on three different mutants, we conclude that L-serine catabolism through the one-carbon metabolism pathway promotes chronological longevity/stationary phase survival under non-restricted conditions.

**Figure 7.**
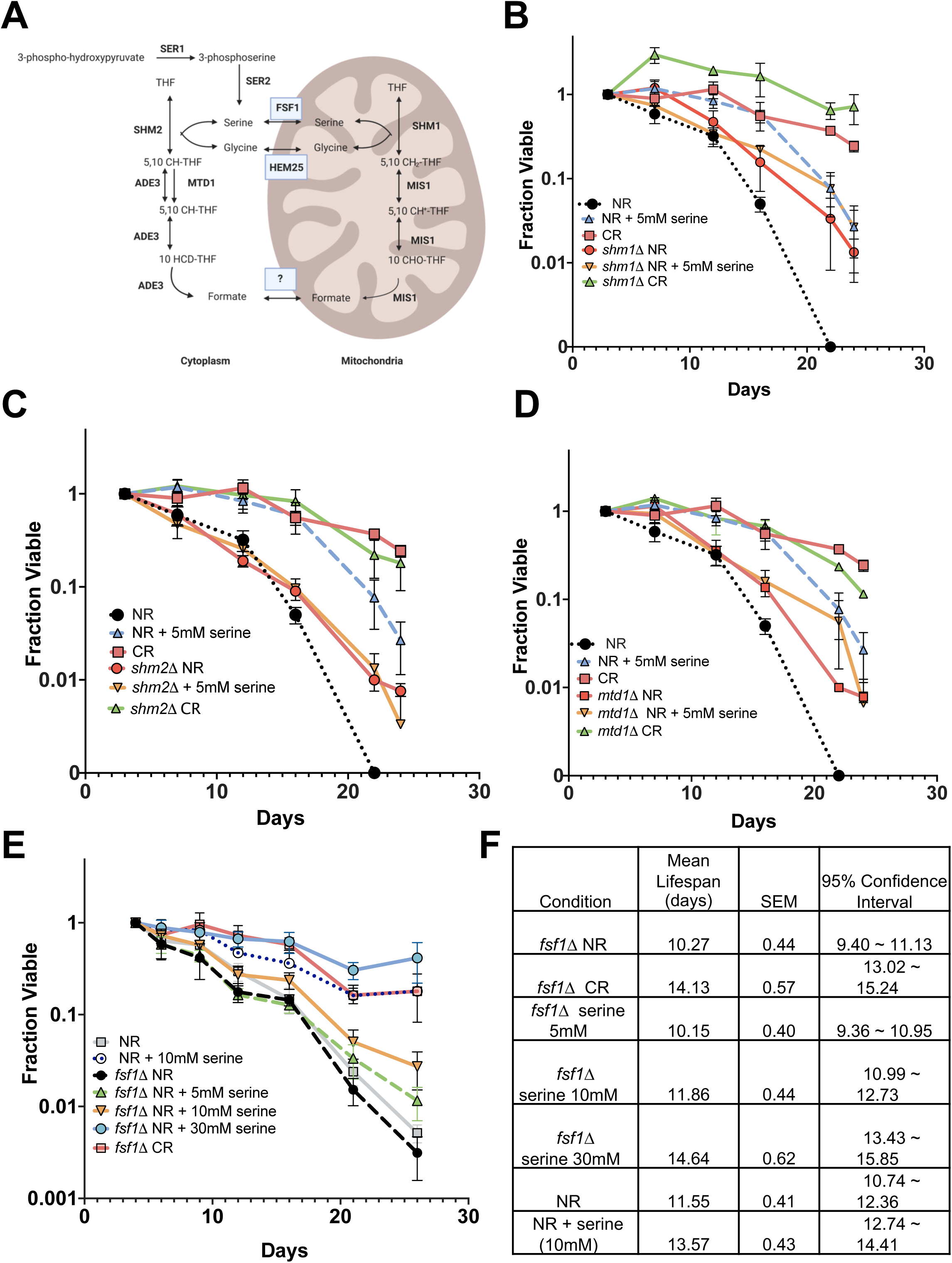
L-serine extends CLS through the one-carbon metabolism pathway. **(A)** Schematic diagram of one-carbon metabolism is *Saccharomyces cerevisiae*. Enzymes that catalyze specific reactions are in bold. Transporters are boxed. **(B)** CLS of BY4741 and *shm1Δ* mutant under NR and CR conditions, or supplemented with 5 mM L-serine. **(C)** CLS of BY4741 and *shm2Δ* mutant under NR and CR conditions, or supplemented with 5 mM L-serine. **(D)** CLS of BY4741 and *mtd1Δ* mutant under NR and CR conditions, or supplemented with 5 mM L-serine. **(E)** CLS of BY4741 and *fsf11Δ* mutant under NR and CR conditions, or supplemented with 5 mM L-serine. **(F)** Mean CLS statistics from panel E calculated using OASIS 2.

## DISCUSSION

At the onset of this study we hypothesized that CR may induce the production of one or more longevity factors, perhaps small molecules, peptides, or even proteins, that are released into the growth medium either through secretion from live cells, or breakdown products from dying cells. Chromatography of CRCM and NRCM clearly indicated the longevity factors were water soluble small molecules, but we were surprised to find the major differences between the two types of conditioned media were amino acids. Several unannotated compounds were more enriched in the CRCM, so at this time we cannot rule out the existence of other compounds with weaker effects on longevity.

### Amino acids as extracellular regulators of lifespan

CR in the context of this study consists of glucose restriction. However, dietary composition, not just overall caloric reduction, plays a critical role in modulating lifespan in multicellular organisms. In *Drosophila*, for example, lower dietary concentrations of yeast (amino acid source) or sugar generally improve lifespan, but moderate concentrations in combination are more optimal (55). Most cells in *Drosophila* or other multicellular organisms are not directly exposed to the environment, so they rely on specialized nutrient ‘sensing’ cells that relay messages about nutrient availability, typically in the form of hormones (56).

Unicellular organisms, on the other hand, must directly respond to nutrient fluctuations in the environment, making them dependent on rapid detection and response to changes in nutrients. Yeast cells have multiple amino acid permeases that are under tight transcriptional and translational control, in order to properly regulate uptake (57). For example, when amino acids are scarce, translation of the *GCN4* mRNA is derepressed. Since Gcn4 is a transcriptional activator for these genes (58), this leads to transcriptional induction of most genes encoding amino acid biosynthetic enzymes, a regulatory process known as general amino acid control (GAAC) (59). The GAAC pathway also integrates with the TOR signaling pathway, which senses nitrogen availability (60), and links amino acid availability to lifespan regulation (Powers 2006 ref?). Activation of GAAC generally reduces CLS, while suppression of GAAC extends CLS (61). This fits well with our finding that supplementing non-restricted cultures with CRCM, which contains abundant amino acids, extends CLS, while NRCM that is amino acid depleted, does not.

Common laboratory yeast strains such as W303, YPH499, and BY4741/BY4742 have several amino acid auxotrophies due to mutations in genes like *HIS3*, *LYS2*, *LEU2*, *TRP1*, or *MET15*. Media containing standard concentrations of auxotrophy-complementing amino acids reduces the final biomass of cultures and shortens chronological lifespan, while excess amounts of these amino acids abrogates the aging phenotype (62). Amino acid uptake has also been genetically implicated in regulation of chronological aging. Chronological lifespan QTL analysis of outbred strains from a cross between S288C and a vineyard yeast strain revealed a polymorphism in the *BUL2* gene (63). *BUL2* encodes a subunit of an E3 ubiquitin ligase that controls trafficking of high affinity amino acid permeases to the vacuole for degradation (64). Reduction in Bul2 function therefore stabilizes the permeases and increases intracellular amino acids, thus increasing TOR activity and shortening CLS. Most recently, availability of non-essential amino acids in the growth medium was shown to be important for chronological longevity (49). Specific amino acids were not critical, but rather the total amount of amino acids functioned to prevent hyperacidification of the growth medium. This scenario could also be at play with the numerous amino acids enriched in CRCM. In the case of L-serine we found that it was capable of buffering pH at higher concentrations, but its catabolism was important at lower concentrations.

Specific amino acids have significant impact on lifespan as well. Branched chain amino acid (BCAA) supplementation has been shown to extend CLS of *S. cerevisiae* (61), and *C. elegans* (65), consistent with the apparent biosynthesis of BCCA we observed under the CR condition. However, BCAA restriction improves late-life health span and lifespan in *Drosophila* and mice (66, 67). Given such large differences in effects between species, this could reflect changes in amino acid balance, rather than direct effects due to BCAA levels (66, 67). Methionine restriction also extends lifespan in all model organisms tested thus far (68). It should be noted that BY4741 is auxotrophic for methionine due to the *met15Δ* mutation, indicating that this strain is already relatively long-lived compared to a strain that is *MET15*^+^ (23). Even with the *met15Δ* mutation, L-methionine or L-cysteine supplementation had little effect on CLS (Figure 5C). Furthermore, since CRCM and L-serine both extended CLS of FY4, the *met15Δ* mutation does not appear to be a major determinant for this cell extrinsic mechanism of lifespan regulation.

Less attention has been placed on L-serine within the aging research community. In addition to our work here and others showing that L-serine supplementation extends yeast CLS (49), L-serine was among the best amino acids at extending *C. elegans* lifespan when supplemented to the worms in a dose dependent manner (69). L-serine supplementation to mice was also recently shown to reduce food intake, improve oxidative stress, and SIRT1 signaling in the hypothalamus of aging mice, though lifespan was not tested (70). Lastly, L-serine is also being studied as a possible neuroprotectant in the treatment of ALS and other neurodegenerative disorders (71–73). Despite these beneficial effects, supplementing with L-serine was reported to be pro-aging when the only other amino acids added to the media were those covering the auxotrophies (74). As with branched chain amino acids, these discrepancies could be due to the combination of auxotrophies and media content, which has been shown to be a major variable driving different CLS results from different labs (35, 39).

### Why do amino acids accumulate in the conditioned media of CR cultures?

In the presence of sufficient glucose, *S. cerevisiae* cells actively suppress respiratory metabolism and biomass production through the TCA cycle, a phenomenon known as the Crabtree effect in yeast, and the Warburg effect in cancer cells (75). When glucose becomes limiting, however, *S. cerevisiae* cells utilize oxidative metabolism over fermentative metabolism, resulting in elevated respiration and electron transport. Under such conditions, amino acids may be used to replenish TCA cycle intermediates through trans- and deamination reactions, a process called anaplerosis. Normally, in cells originally grown in 2% glucose (NR), glucose depletion triggers increased amino acid uptake that involves upregulation of permeases via TOR (76). Initial growth under CR (0.5% glucose) appears to generally reduce amino acid uptake as indicated by accumulation of amino acids we observe in the conditioned medium (Figure 4A-D), and instead prioritizes consumption of alternative carbon sources such as acetate (Figure 6A; (37, 38)), which yeast cells can convert into acetyl-CoA for TCA intermediate replenishment, or gluconeogenesis via the glyoxylate cycle (77). This likely better accommodates the increased storage of glycogen and trehalose induced by CR and associated with long term cell survival in stationary phase (20). The higher cell densities (biomass) achieved by NR cultures instead places tremendous demand for synthesis of macromolecules associated with cell growth, such as nucleotides, lipids, and proteins, thus depleting amino acids from the media.

CR could also potentially make ammonium sulfate a preferred nitrogen source over the amino acids that are usually preferred under the non-restricted conditions. Ammonium sulfate has been shown to reduce CLS and is actually left out of the custom HL medium designed to optimize CLS (48, 78). Therefore, assimilation of the ammonium under CR could potentially extend CLS by reducing ammonium toxicity, similar to the CR-induced consumption of acetic acid (37, 38). Evidence for this mechanism comes from studies of amino acid preference during fermentation by wine yeasts (79). Of the 17 amino acids tracked, lysine was utilized the fastest, followed by a group of 10 (Asp, Thr, Glu, Leu, His, Met, Ile, Ser, Gln, Phe) that were consumed quicker than ammonium sulfate, and 6 (Val, Arg, Ala, Trp, Tyr, Gly) that were slower. Consistent with this hypothesis, lysine was the most depleted amino acid in BY4741 NR conditioned media, and was partially rescued by CR (Figure 4A and B). Moreover, all of the fast-depleted amino acids in wine fermentation, except glutamine, were depleted under NR and rescued by CR (Figure 4A and B).

### One-carbon metabolism in regulation of aging

Although multiple amino acids are more abundant in CRCM than NRCM, we focused on L-serine because the biosynthesis gene *SER1* is a QTL for CLS in the BY4741 background (46). L-serine is also a major entry point for the one-carbon metabolism pathway, and a key component of the transsulfuration pathway (80), which has been implicated in longevity (81). The one-carbon metabolism pathway supports multiple cellular processes such as biosynthesis of purines, amino acid homeostasis (glycine, serine, and methionine), epigenetics through SAM and chromatin methylation, and redox defense (82). However, few studies directly linked it to the regulation of aging. In one study, activation of naïve T cells from aged mice was attenuated because of a deficit in the induction of one-carbon metabolism enzymes (83). In our current study, L-serine supplementation extended CLS in a manner dependent on the one-carbon metabolism pathway (Figure 7), which we interpret as the one-carbon units donated from L-serine allowing cells to complete biosynthesis processes required to effectively enter quiescence. Of note, L-glycine was not depleted from NR yeast cultures and had no effect on CLS when supplemented (Figures 4A and 5C). In this sense, non-restricted yeast cells may be similar to cancer cells that rely on exogenous serine, but not glycine, for proliferation (84). Overexpression of serine hydroxymethyltransferase SHMT2 is also associated with poor prognosis in cancer patients, while downregulating this enzyme suppresses tumorigenesis in human hepatocellular carcinoma (85).

Yeast cells lacking the mitochondrial serine hydroxymethyltransferase Shm1, or the NAD-dependent 5,10-methylenetetrahydrofolate dehydrogenase Mtd1, each displayed extended mean and maximum CLS. Cells lacking the cytoplasmic Shm2 enzyme also appeared to extend maximum CLS (Figure 7). These results suggest that perturbing flux through the one-carbon metabolism pathway under non-restricted conditions can influence long term cell survival, perhaps by forcing metabolism toward gluconeogenesis and the glyoxylate cycle. Interestingly, yeast replicative lifespan extension caused by deletion of the *RPL22A* gene was recently shown to correlate with reduced translation of one-carbon metabolism enzymes (86). Furthermore, deleting genes involved in one-carbon metabolism moderately extended replicative lifespan, similar to what we observed for CLS. We therefore hypothesize that CR may also reduce flux through the one-carbon metabolism pathway, consistent with reduced serine consumption from the media and strong lifespan extension of the *shm1Δ*, *shm2Δ*, or *mtd1Δ* mutants by CR. The impact of these perturbations, which represent evolutionarily conserved and highly connected pathways, may depend on genetic and environmental context, and thus the yeast model is ideal for further systematic experimental characterization (87). Given the common effect of one-carbon metabolism on yeast RLS and CLS, and its strong conservation from yeast to mammals, future investigation of its roles in metazoan aging models is warranted.

## MATERIALS AND METHODS

### Yeast strains and media

*S. cerevisiae* strains used in this study were BY4741 (*MAT***a** *his3Δ1 leu2Δ0 met15Δ0 ura3Δ0)*, FY4 (*MAT***a** prototrophic), and several deletion mutants from the Euroscarf yeast knockout (YKO) collection (88). Synthetic complete (SC) growth medium was used for all experiments except for the use of custom ‘human-like’ HL media (35, 48). Glucose was added to a final concentration of either 2.0% (non-restricted [NR]) or 0.5% (calorie restricted [CR]). To supplement amino acids, SC or HL medium was used containing 2% glucose (NR), and with amino acids added to a final concentration of 5 mM, 10 mM, 20 mM or 30 mM where indicated. Unless noted otherwise, all cultures (10 ml) were grown at 30°C in 15 ml glass culture tubes with loose fitting metal caps on a New Brunswick Scientific roller drum.

### Semiquantitative (spot) and quantitative assays for measuring CLS

To assess chronological lifespan, overnight 10 ml SC 2% glucose (NR) cultures were started from single colonies in triplicate. Next, 200 µl of the overnight cultures was used to inoculate fresh 10 ml cultures of the indicated SC medium conditions (NR, CR, or NR + amino acid). For each time point, 20 µl aliquots were then removed from cultures at the indicated times in stationary phase (starting at day 3) and serially diluted in 10-fold increments with sterile water in 96-well plates. For semi-quantitative spot assays, 2.5 µl of each dilution was spotted onto YPD 2% glucose agar plates as previously described (6). The plates were then digitally imaged on a gel documentation system after 2 days of colony growth and the time points were compiled together to visualize the changes in viability over time. For the quantitative CLS assays, 2.5 µl of the 1:10, 1:100, and 1:1,000 dilutions of each culture were spotted onto YPD plates that were then incubated at 30°C for 18 to 24 h to allow for microcolony formation (16). We typically spot the three dilutions for 12 different cultures onto one YPD plate. Images of each dilution spot were captured on a Nikon Eclipse E400 tetrad dissection microscope at 30x magnification such that the entire spot fills the field of view. Microcolonies were then counted from the images either manually with a counting pen, or automatically using OrganoSeg, a program originally developed for counting mammalian organoids in culture (89), which we have adapted for counting yeast colonies (Enriquez-Hesles et al., manuscript in preparation). At the end of each experiment, percent viability was calculated for each time point by normalizing to the first day of colony forming unit (CFU) measurements. Standard deviation error bars on the survival curve graphs were determined from 3 biological replicates. Statistical analysis was performed using the online program OASIS 2 (90), reporting mean life span (days), standard error of the mean (SEM), and 95th percentile confidence interval (95% CI) of the mean for each strain or condition.

### Preparation of conditioned media concentrates

To collect and concentrate conditioned media for CLS assays and amino acid profiling, BY4741 or FY4 strains were grown in 150 mL SC cultures (with either 2% glucose or 0.5% glucose) at 30°C for 5 days in a shaking water bath. The cultures were then centrifuged and the supernatants were condensed from 150 mL down to 15 mL using a Büchi Rotavapor-R apparatus, then filtered by passing through a 0.2 micron filter and stored at −20°C. For supplementation experiments, conditioned media concentrates (derived from NR or CR cultures) were added to 10 mL of non-restricted SC-NR media to final concentrations of 0.5%, 1%, or 2% where indicated.

### Chromatography

Fifteen BY4741 NR or CR cultures (10 ml SC each), were grown to saturation for 5 days at 30°C in 18 mm glass test tubes with loose fitting metal caps. The glassware was acid washed with 0.1N HCl before use. Cells were pelleted by centrifugation and the conditioned media pooled. The pooled media was then concentrated in a Büchi Rotavap from 150ml down to approximately 2.5 ml. The concentrates were centrifuged at 4,000 rpm (2,987 RCF) for 10 minutes in 15 ml conical tubes to remove any solid precipitates. Two ml of the clarified concentrate was loaded onto a 1 x 26 cm Sephadex G-10 column, and fractionated with double distilled water. 2 ml fractions were collected by gravity flow in a Pharmacia fraction collector and filter sterilized through 0.22µm syringe filters. The fractions were then added to new 5 ml CLS cultures at a 1:5 ratio (ml concentrate: ml culture) and semiquantitative CLS assays performed. NaCl (100 mM) was eluted through the column before and after the media concentrates to determine the gel size retention fractions as measured by electrical conductivity.

### Metabolomics

BY4741 NR and CR cultures (10 ml SC each) were grown to stationary phase (day 5), then centrifuged in 15 ml disposable conical tubes (Falcon). The supernatant media was filter sterilized through 0.22µm syringe filters and frozen at −80°C. Untargeted metabolomics of conditioned media from 6 NR and 6 CR cultures was performed via gas chromatography/electron-ionization mass spectrometry (GC/ei-MS) in the Metabolomics Laboratory of the Duke Molecular Physiology Institute (DMPI), as described (91). Metabolites were extracted by the addition of methanol. Dried extracts were methoximated, trimethylsilylated, and run on an 7890B GC-5977B ei-MS (Agilent Corporation, Santa Clara, CA), with the MS set to scan broadly from *m/z* 50 to 600 during a GC heat ramp spanning 60° to 325 °C. Deconvoluted spectra were annotated as metabolites using an orthogonal approach that incorporates both retention time (RT) from GC and the fragmentation pattern observed in MS. Peak annotation was based primarily on DMPI’s own RT-locked spectral library of metabolites, which is now one of the largest of its kind for GC/EI-MS. DMPI’s library is built upon the Fiehn GC/MS Metabolomics RTL Library (a gift from Agilent, their part number G1676-90000; (92)). Quantities from the mass spectrometry were normalized to OD_600_ of the cultures to account for cell density. 160 metabolites were annotated based on matches with a spectral library. Another 115 metabolites were not matched in the library and remain unannotated (Table S1).

### Quantitative amino acid profiling

Conditioned NR and CR media from day 5 stationary phase cultures was collected and concentrated with the Rotavap as described above. As a control, SC media without glucose was also concentrated and analyzed. Samples were submitted to the UVA Biomolecular Analysis Facility and then analyzed using a ZipChip system from 908 Devices that was interfaced with a Thermo Orbitrap QE HF-X Mass Spectrometer. Samples were prepared by diluting 10 µL with 490 µL of LC-MS grade water, which was then further diluted 1:10 with 90 µl of the ZipChip diluent (908 Devices Inc., P/N 810-00168). The samples were loaded onto ZipChip HR Chip (908 Devices Inc., P/N 810-00194) for analysis. The following ZipChip analysis settings were utilized: Field strength: 500V/cm, Injection volume: 7 nl, Chip Type: HR, BGE: Metabolite, Pressure assist: Enable at 7 minutes, Run time: 10 minutes, MS setting (Thermo Orbitrap QE HF-X), m/z range: 70-500, Resolution: 15000, 1 microscan, AGC target: 3E6, Max ion injection time: 20 ms, Inlet capillary temperature: 200°C, S Lens RF: 50.

### RNA analysis

Cells from non-restricted overnight cultures were inoculated into 75 ml of fresh non-restricted SC medium that was supplemented with 1.5 ml of concentrated conditioned media (CRCM or NRCM) or 1.5 ml of sterile water as a control. The starting OD_600_ was 0.05 in 250 ml Erlenmeyer flasks. Cultures were grown at 30°C in a New Brunswick water bath shaker. For the log phase condition, 50 ml of the samples were collected at OD_600_ of 0.2. Equivalent numbers of cells were collected from smaller aliquots harvested at 24 hr and 96 hr. Total RNA was isolated using the hot acid phenol method and then processed into Illumina DNA sequencing libraries as previously described (50), with slight modifications. Briefly, total RNA was treated with DNase I for 10 min at 37°C and then measured for concentration and quality with an Agilent Bioanalyzer. PolyA mRNA selection was performed on 5 µg of the DNase-treated total RNA with the NEBNext Poly(A) mRNA magnetic isolation module (E7490). DNA sequencing libraries were then generated with the NEBNExt Ultra Directional RNA library Prep kit for Illumina (E7420). Libraries were sequenced on an Illumina NextSeq 500 by the UVA Genome Analysis and Technology Core (GATC). Sequencing files are available at GEO (accession number GSE151185). Sequencing reads were mapped to the sacCer3 genome using bowtie2 with default settings (93). We preprocessed sequencing data from the UVA GATC and analyzed differential gene expression in R using DESeq2 (94).

### Acetic acid measurements

100 µl aliquots were taken at designated time points from standard 10 ml CLS cultures. Cells were pelleted by centrifugation at 2,500 rpm at 4°C, and 50 µl of supernatant was removed and stored at −80°C, until further analysis. Acetic acid concentration for each sample was then later determined using an Acetic Acid Kit (Biopharm AG) per manufacturer’s instructions. The acetic acid concentrations and standard deviations provided are an average of three biological replicates for each condition and reported as g/L.

## ACKNOWLEDGMENTS

We thank Joel Hockensmith for small molecule chromatography advice and Mark Okusa for use of a Rotavap apparatus. We also thank Arun Dutta and Katherine Owsiany for assistance with R and RNA-seq analysis. E.E.H was supported by the Medical Scientist Training Program (MSTP) training grant T32GM007267, and the Cell and Molecular Biology (CMB) training grant T32GM008136 from the NIH. R.D.F was also supported by the CMB training grant. This work was also supported by NIH grants R21AG053596 and RO1GM075240 to J.S.S., R56AG059590 to J.L.H and J.S.S., RO1AG043076 to J.L.H., RO1AG045351 to M.D.H, and RO1CA214718 to K.A.J., who was also supported by The David and Lucile Packard Foundation (#2009-34710).

## SUPPLEMENTAL FIGURE LEGENDS

**Figure S1.**
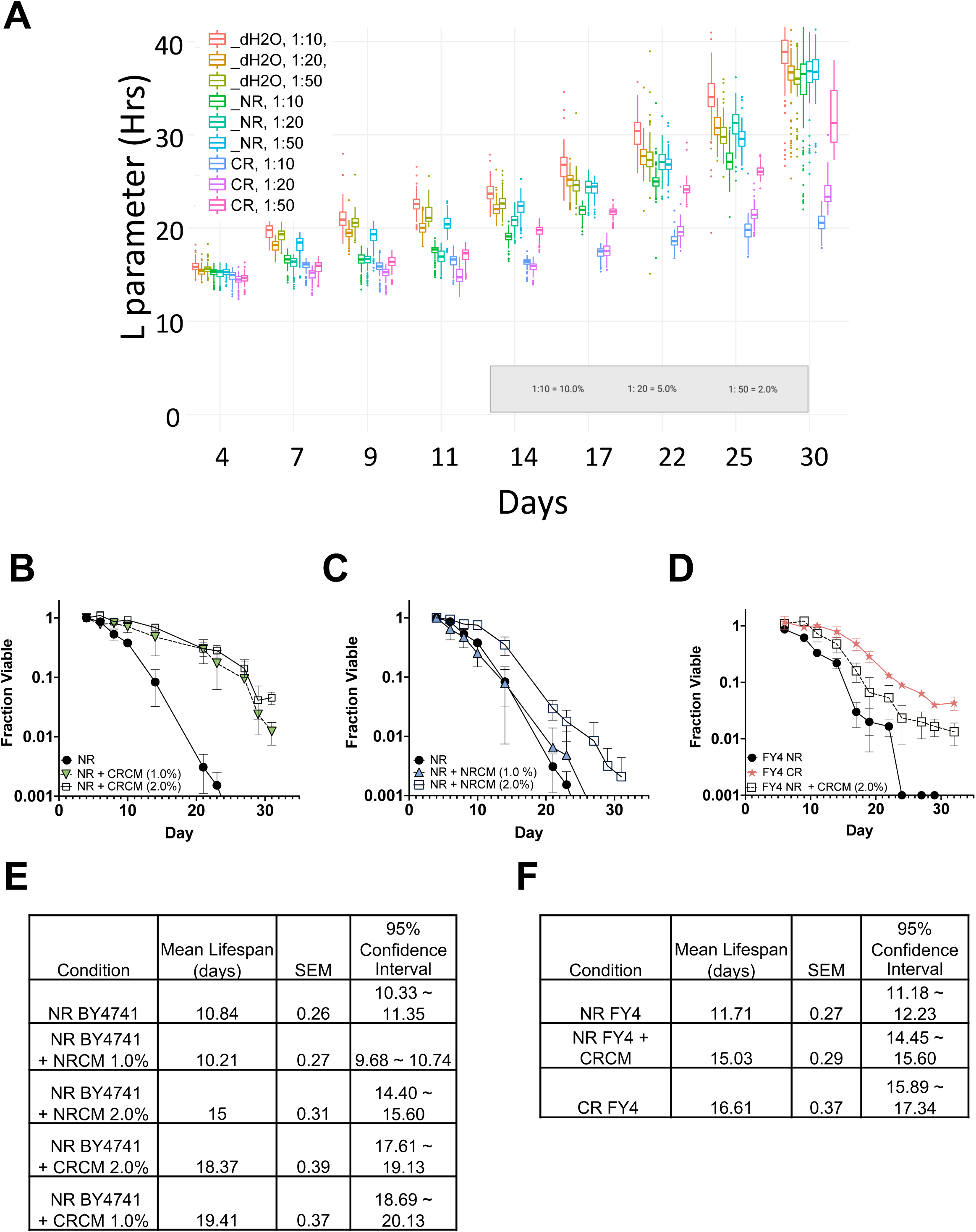
Longevity factors are present in conditioned media from glucose-restricted yeast cultures. **(A)** Quantitative high throughput cell array phenotyping (Q-HTCP) assay for detecting longevity factor activity. BY4741 was inoculated into SC NR (2% glucose) media, in 50 µl volumes in 384-well ‘Aging Arrays’. Cultures were treated with 10X-concentrated conditioned media from 7-day old CR (0.5% glucose) or NR (2% glucose) cultures or treated with an equal volume of water, as indicated in the legend (top left). At the indicated days, cells from the aging arrays were printed onto YPD Growth Array plates, and L values in hours were obtained. A strong dose response to longevity factors in CR conditioned media was detected, as indicated by lower L values. Each box in the plot represents the distribution of 96 replicate cultures. **(B)** Using BY4741, NR CLS cultures were supplemented with the concentrated CRCM to 2.0% or 1.0% (vol/vol). **(C)** Using BY4741, NR CLS cultures condition were supplemented with concentrated NRCM to 2.0% or 1% (vol/vol). Panels B and C have the same NR control. **(D)** CLS of non-restricted (NR) FY4 supplemented at 2% (vol/vol) with CRCM derived from BY4741. FY4 was also grown under the CR condition as a control. **(E)** Mean CLS statistics from panels B and C, calculated using OASIS 2. **(F)** Mean CLS statistics from panel F calculated using OASIS 2.

**Figure S2.**
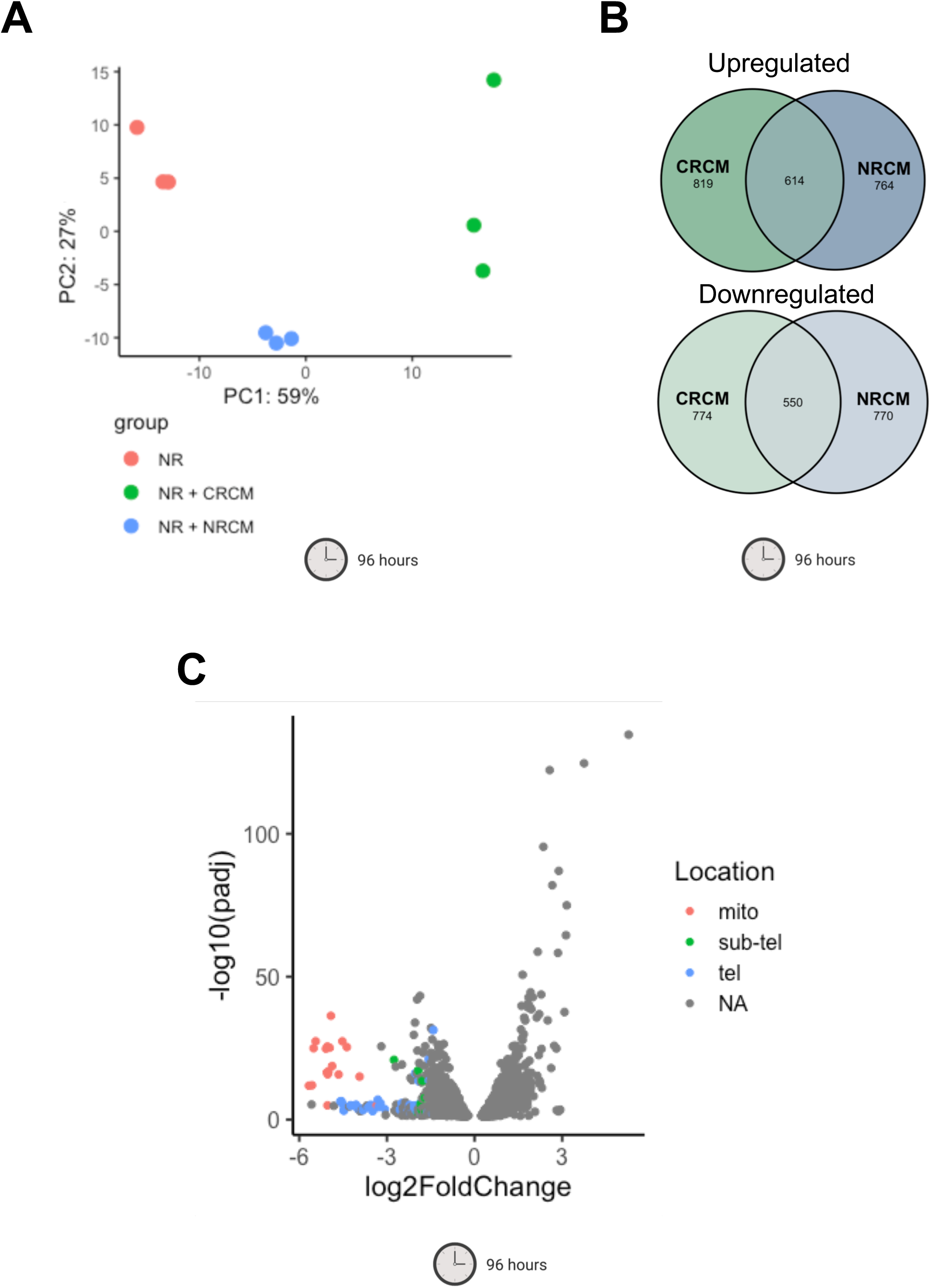
RNA-seq analysis of gene expression changes induced by CRCM and NRCM. **(A)** Principal component analysis of RNA-seq samples collected at 96 hours. **(B)** Venn diagram of differentially expressed genes (up or down; FDR<0.05) for NRCM or and CRCM compared to the NR control at 96 hrs. **(C)** Volcano plot displaying differential expressed genes compared between NR + CRCM and NR control samples at 96 hrs. The vertical axis (y-axis) corresponds to the p-adjusted value and the horizontal axis (x-axis) displays the log2-fold change value. Red, green, and blue denotes genes located in the mitochondrial genome, sub-telomeric, or telomeric regions, respectively.

**Figure S3.**
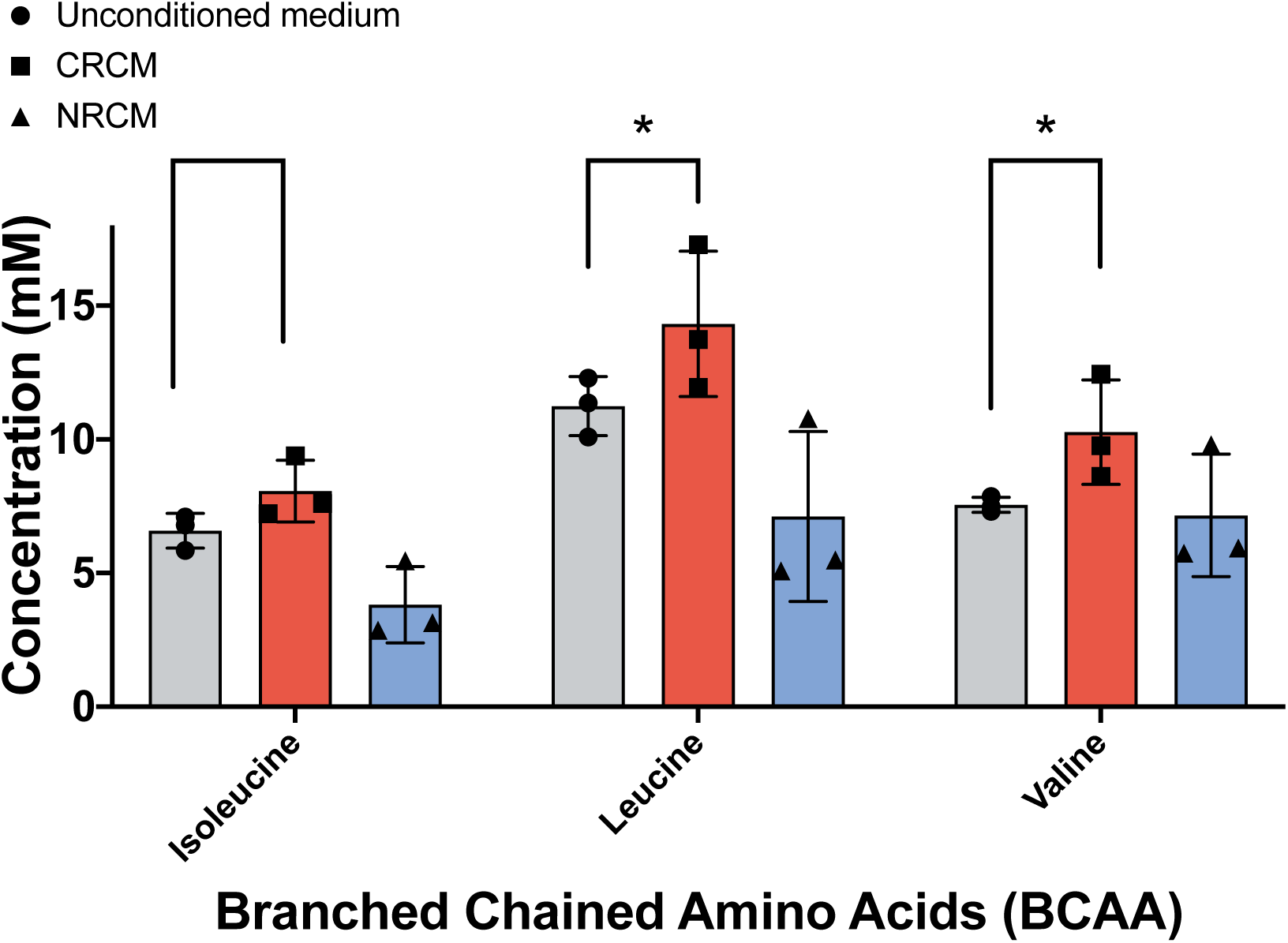
Branched chain amino levels in concentrated CRCM or NRCM. L-isoleucine, L-leucine, and L-valine levels were measured in concentrates of CRCM, NRCM, or unconditioned SC media. Significant increases in CRCM over the starting SC media are indicated by an asterisk (p<0.05, student’s t-test).

**Figure S4.**
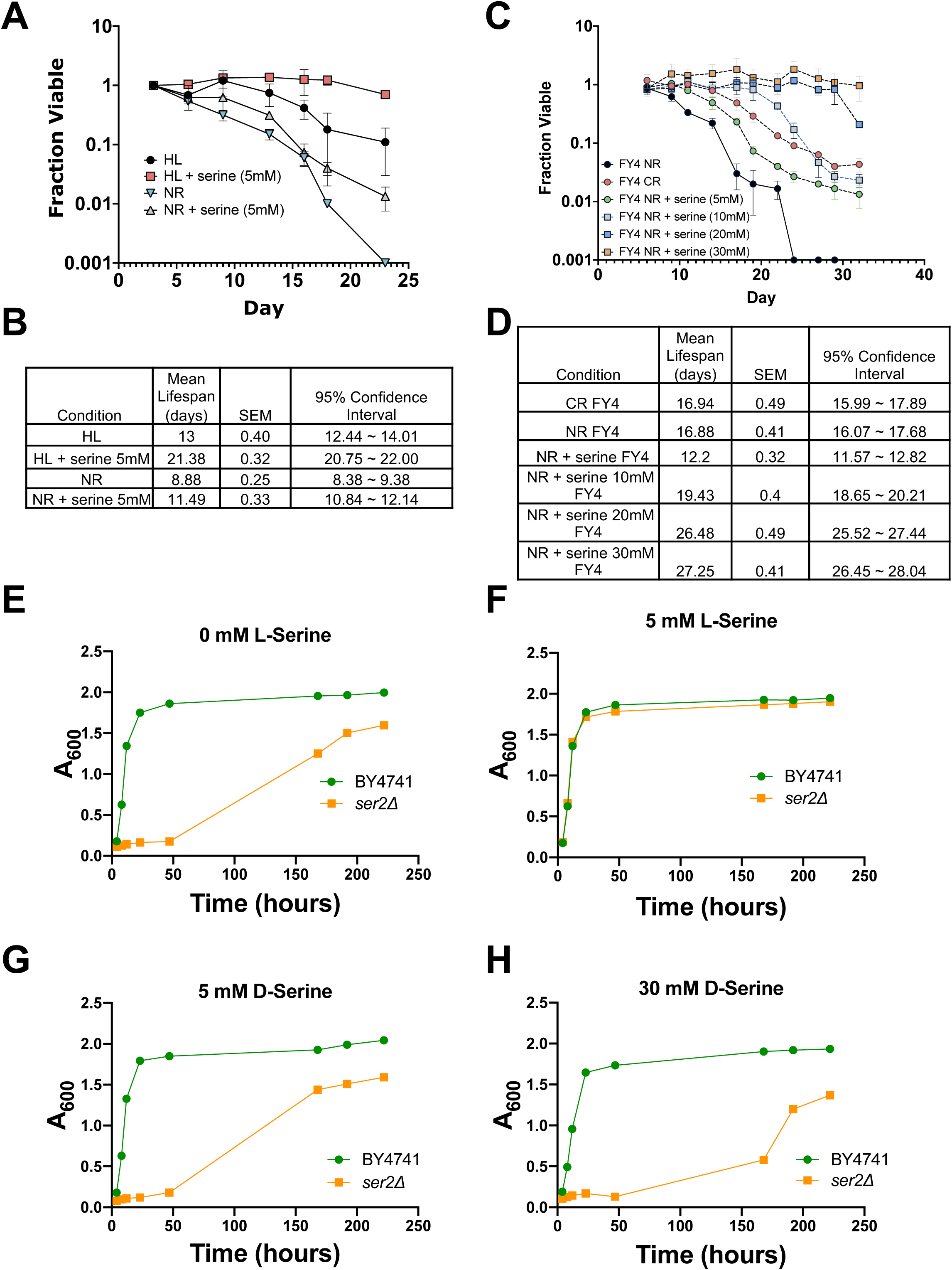
Analysis of L-serine and D-serine effects on CLS and cell growth. **(A)** CLS of BY4741 cells grown in SC (NR) media or custom Human-Like (HL) medium supplemented with 5 mM L-serine. **(B)** Mean CLS statistics from panel A, calculated using OASIS 2. **(C)** CLS of prototrophic strain FY4 in SC (NR) media supplemented with L-serine at the indicated concentrations. FY4 was also grown in SC (CR) media as a control. **(D)** Mean CLS statistics from panel C, calculated using OASIS 2. **(E-H)** Growth curves of BY4741 and *ser2Δ* mutant in SC-serine media supplemented with the indicated concentrations of L-serine or D-serine (mean, n=2).

**Figure S5.**
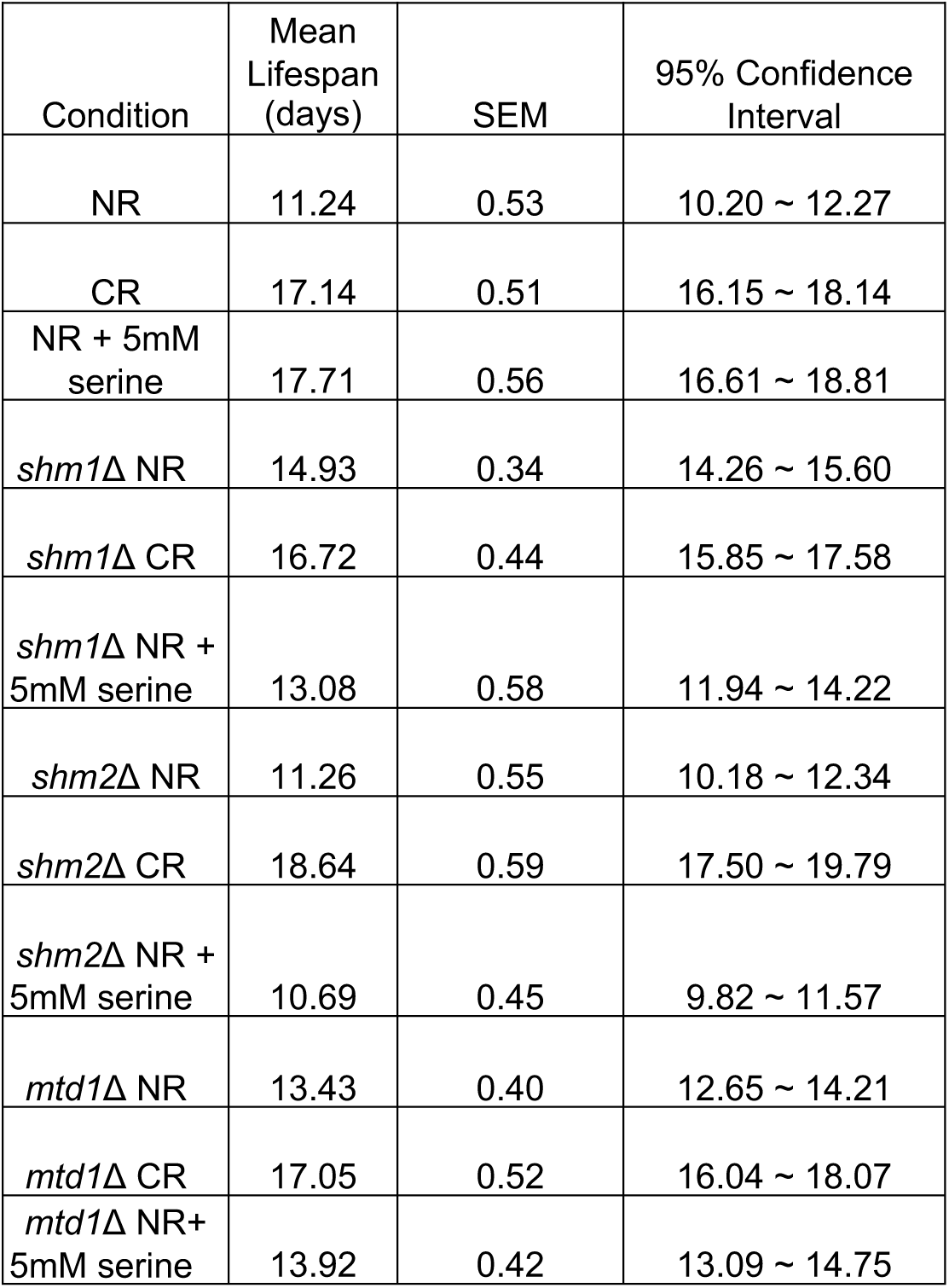
Statistical analysis of CLS assays for one-carbon metabolism deletion mutants. Statistics for CLS assays presented in Figure 7, panels B, C, and D. The mean CLS, SEMs, and 95% confidence intervals were calculated using OASIS 2.

